# Genomic footprints of selfing, introduction history, and long-distance dispersal in an invasive alien plant

**DOI:** 10.64898/2026.05.03.722469

**Authors:** Raúl Sánchez-García, Andy J. Green, María A. Ortiz, Cristina García, Francisco Hortas, Chevonne Reynolds, Jennifer Rowntree, Ester A. Serrão, Lina Tomasson, Karin Tremetsberger, Casper H.A. van Leeuwen, Joaquín Ortego

**Affiliations:** Department of Conservation Biology and Global Change, Doñana Biological Station (EBD), CSIC, Seville, Spain; Department of Natural Sciences, Manchester Metropolitan University, Manchester, UK; Department of Plant Biology and Ecology, University of Sevilla, Seville, Spain; Department of Biological Sciences, School of Life and Environmental Sciences, Royal Holloway University of London, Egham, Surrey, UK; Department of Biology, Institute of Marine Research (INMAR), University of Cadiz, Puerto Real, Spain; School of Animal, Plant and Environmental Sciences, University of the Witwatersrand, Johannesburg, South Africa; School of Biological and Marine Sciences, University of Plymouth, Drake Circus, Plymouth, UK; Centre of Marine Sciences, CCMAR, University of Algarve, Campus de Gambelas, Faro, Portugal; National Coordinator for Aquatic Invasive Alien Species, Swedish Agency for Marine and Water Management, Gothenburg, Sweden; BOKU University, Institute of Botany, Department of Ecosystem Management, Climate and Biodiversity, Vienna, Austria; Department of Ecology, Radboud Institute for Biological and Environmental Sciences, Radboud University, Nijmegen, the Netherlands; Department of Ecology and Evolution, Doñana Biological Station (EBD), CSIC, Seville, Spain

**Keywords:** admixture, buttonweed, *Cotula coronopifolia*, genetic diversity, genomic of invasions, outcrossing, seed dispersal, selfing

## Abstract

Biological invasions are natural experiments for studying the evolutionary and ecological processes underlying colonization success and range expansion. Using genome-wide data — generated via genotyping-by-sequencing (GBS) from 30 populations spanning Europe and South Africa— we investigated the colonization history and successful spread of the invasive buttonweed *Cotula coronopifolia*, an annual plant introduced into Europe from South Africa about 300 years ago. Our analyses identified three major lineages in Europe distributed across the continent, often co-occurring without evidence of admixture. Phylogenomic dating revealed that these lineages diverged > 2,000 years ago —well before the earliest European records— suggesting divergence within the native range and either multiple introductions or a single introduction with multiple lineages. Mating-system inference shows that reproduction occurs primarily via self-fertilization (∼70% on average), although outcrossing predominates in some populations, revealing a facultative mating system. This high selfing rate has led to extremely low heterozygosity in most populations and a strong genetic structure. Genetic clustering also revealed admixed individuals resulting from rare inter-lineage outcrossing; comparisons of empirical and simulated data indicate that increased diversity after sporadic admixture events decays rapidly under subsequent selfing. Both the introduction history and long-distance dispersal facilitated by waterbirds likely explain the scattered distribution of lineages across Europe. Altogether, these results provide an empirical demonstration of Baker’s “ideal weed” concept, highlighting the role of a flexible mating system in providing reproductive assurance during colonization and showing how predominant selfing shapes the genomic landscape of an invasive species.

**SIGNIFICANCE STATEMENT:** Understanding how reproductive strategies influence colonization and spread of alien species is central to invasion biology. By combining population genomics and phylogenomic inference, this study provides key insights into the colonization history and successful invasion of the self-fertilizing plant *C. coronopifolia*, introduced from South Africa to Europe in the 18^th^ century. We show that the species’ invasion success relies on the introduction of multiple lineages and the predominance of selfing, which has drastically reduced genetic diversity yet contributed to reproductive assurance and spread across diverse habitats. Occasional outcrossing and long-distance dispersal by waterbirds or through horticultural transport have further shaped the species’ genetic landscape. These findings illustrate how self-compatibility and ecological generalism can overcome genetic constraints during range expansion and provide the basis for understanding the evolutionary dynamics of selfing plant invasions.

## INTRODUCTION

Biological invasions have long captured scientific attention, as they often pose a direct threat to biodiversity and the conservation of ecosystems (Simberloff et al. 2013; Pyšek et al. 2020). In addition to their ecological and socioeconomic impacts, invasions also serve as natural experiments in evolution, providing opportunities to study how species reach distant locations, spread and establish outside their native range, and adapt to new environments (Bock et al. 2015). Advances in next-generation sequencing (NGS) technologies have further expanded the scope of invasion genetics, allowing us to uncover the demographic and evolutionary processes underpinning invasion success across diverse taxa (Rius et al. 2015).

A common assumption in invasion biology is that high genetic variability is critical for adaptive success, as it provides the raw material for evolution and enables populations to respond to selective pressures in new environments (Dlugosch et al. 2015). However, recently established populations of invasive species often experience reductions in genetic diversity due to founder effects and demographic bottlenecks, creating an apparent contradiction between theoretical expectations and empirical observations (Schrieber and Lachmuth 2017). This inconsistency is known as the “genetic paradox of invasion” (Allendorf and Lundquist 2003). Several non-exclusive mechanisms have been proposed to resolve this paradox, including phenotypic plasticity, multiple introductions leading to admixture, self-fertilization, clonal or asexual reproduction, and hybridization (Liu et al. 2006; Roman and Darling 2007; Hao et al. 2011; Rius and Darling 2014; Williams et al. 2014). One scenario in which the genetic paradox is particularly relevant is described by Baker’s law, which states that colonization success of remote regions is more likely in self-compatible organisms, as a single propagule would be capable of establishing a sexually reproducing colony (Baker 1955). While this strategy facilitates colonization, it may also constrain genetic diversity, creating a potential trade-off between short-term establishment success and long-term evolutionary potential (Pannell et al. 2015). In plants, the interplay between self-compatibility, clonal reproduction, and phenotypic plasticity can therefore strongly influence invasion outcomes (Rollins et al. 2013).

The buttonweed *Cotula coronopifolia* (Asteraceae) provides an excellent model system to explore these processes. This annual or short-lived perennial herb inhabits a variety of wet to seasonally wet habitats, ranging from freshwater to brackish wetlands. It exhibits dual reproductive strategies: asexual vegetative propagation via rooting nodes and sexual outcrossing or self-fertilization through hermaphroditic florets (van der Toorn 1980). Originally native to South Africa, *C. coronopifolia* has spread globally and is now established across Europe, the Americas, Australia, and other regions (van der Toorn 1980; Marfella et al. 2023) (Fig. 1). In Europe, the species was first recorded in Emden, Germany (∼1739), followed by the Netherlands (∼1742), Sweden (∼1853), and later Spain and the United Kingdom (∼1886) (Ridley 1930; van der Toorn 1980; Wilkinson 2023). The species can exclude native plants and continues to expand in Europe, along the Baltic coast and into newly created wetlands (Tomasson 2020). Early introductions are thought to have occurred via shipping, particularly through ballast water transport between South Africa and Europe, and possibly through seed transport in wool (van der Toorn 1980; Verloove and Gonggrijp, 2025). It is now sold in Europe as a pond plant and may potentially escape from gardens (Wilkinson 2023). However, its current distribution, spanning both coastal and inland wetlands, is difficult to explain solely by human-mediated dispersal. Seeds can disperse on water, but field observations have long suggested that waterbirds may contribute to its active dispersal (van der Toorn 1980). The role of birds as competent dispersal vectors has now been empirically demonstrated, with studies confirming that *C. coronopifolia* seeds survive gut passage and that their germination is enhanced thereafter, which has important implications for understanding the specieś invasion ecology and spread in its introduced range (Raulings et al. 2011; Lovas-Kiss et al. 2019; Sánchez-García et al. 2024).

**Fig. 1.**
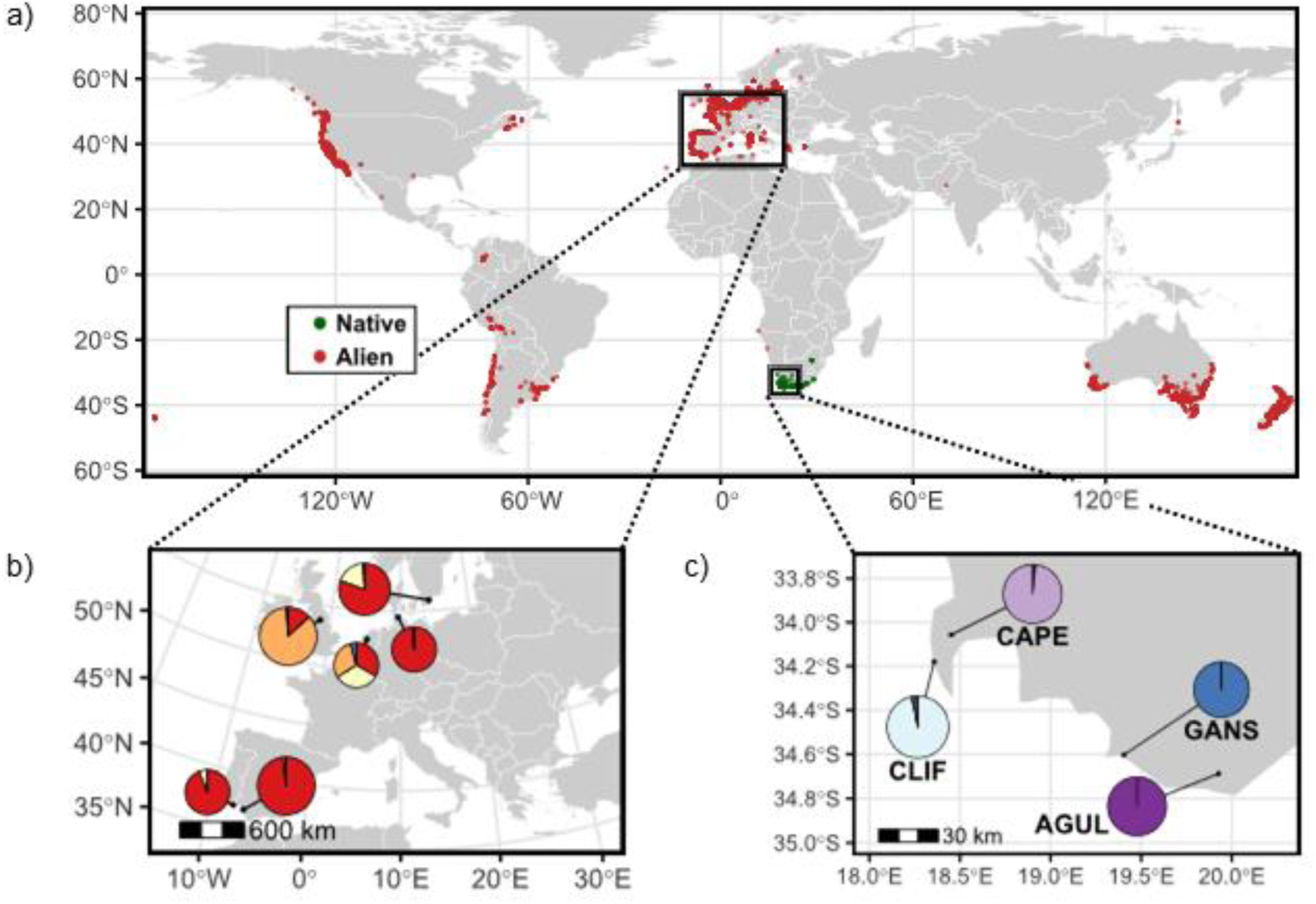
(a) Occurrence records (18,646 observations) of *C. coronopifolia* in its native (green) and alien (red) ranges (Global Biodiversity Information Facility, GBIF, 2015–2025). Records were retrieved via the *rgbif* R package; occurrences within South Africa were classified as native, whereas all others were classified as alien. Only unique, georeferenced presence records passing basic quality filters are shown. Panels (b) and (c) show the results of genetic assignments (pie charts) based on the Bayesian clustering analyses implemented in STRUCTURE for (b) European (5,693 SNPs) and (c) South African (13,079 SNPs) populations. For European populations, pie charts represent the mean probability of assignment to each genetic cluster for samples from each country.

Several characteristics of *C. coronopifolia* align directly with mechanisms proposed to resolve the genetic paradox of invasion. Its capacity for self-fertilization fits the predictions of Baker’s law, enabling establishment from single propagules in novel habitats, while vegetative propagation ensures the persistence and local spread of successful genotypes even in the absence of high genetic diversity. These traits are consistent with Baker’s (1965) concept of the “ideal weed” and the “general-purpose genotype”, which emphasize broad ecological tolerance, reproductive assurance, and the ability to survive under a wide range of environmental conditions. In *C. coronopifolia*, tolerance to variable hydrological and salinity regimes suggests that phenotypic plasticity contributes to its invasive potential (Costa et al. 2009; Marfella et al. 2023). Finally, its capacity for long-distance dispersal by waterbirds (Lovas-Kiss et al. 2019; Sánchez-García et al. 2024) increases the likelihood of colonization events, reinforcing its status as a species with a “general-purpose genotype” capable of colonizing and persisting in diverse habitats.

Despite its nearly cosmopolitan distribution and early introduction into Europe, surprisingly little is known about the invasion history of *C. coronopifolia*. Most previous research has focused on its ecology and morphology (e.g., Costa et al. 2009; Marfella et al. 2023), while key genetic and demographic aspects underlying its invasive success remain unexplored. To fill these gaps, we sampled and obtained genome-wide data from populations of *C. coronopifolia* across both its native range in South Africa and its invasive range in Europe. Specifically, we address three global questions: (i) what does genomic evidence reveal about the colonization history of *C. coronopifolia* in Europe, particularly regarding the number, origin, and timing of introduction events that shaped its current genetic makeup?; (ii) To what extent might reproductive traits traditionally associated with Baker’s “ideal weed” concept –specifically self-compatibility and selfing rate– have contributed to the establishment and successful expansion of the species in its invasive range?; and (iii) what genetic footprints have colonization history, spread by secondary dispersal, and reproductive mode left on European populations? By integrating phylogenomic inference, demographic reconstructions, estimates of selfing rates, and quantification of population genetic structure, diversity and admixture, our study provides a comprehensive population-genomic assessment of the colonization and spread of a facultatively selfing species, contributing to the open debate on how reproductive assurance and ecological generalism interact to shape invasion success despite extraordinarily low levels of genetic diversity.

## RESULTS

### Genomic datasets

We obtained genotyping by sequencing (GBS) data from 266 plants collected at 30 locations. The average number of reads per individual was 9,253,942 (range = 528,852-34,055,893; median = 10,530,730). After the different filtering steps in IPYRAD v. 0.9.93 (Eaton and Overcast 2020), the dataset including all genotyped individuals contained 6,661 unlinked SNPs represented in >90% of the individuals and an average proportion of missing data per individual of 6.5% (range = 0.6-67.4%, median = 1.59%). For the dataset with European populations, we obtained a total of 5,692 of unlinked SNPs with an average of 6.5% missing data (range = 0.6-66.3%; median = 1.32%), while the dataset only including South African populations contained 13,079 unlinked SNPs with 3.98% of missing data (range = 0.9-11.6%; median = 2.79%). The number of SNPs differed between the dataset containing only populations from Europe and the one including only plants from South Africa, owing to the higher genetic diversity observed in the latter (Table 1; Wilcoxon rank-sum test, *W* = 940; *n* = 230, 36; *p* < 0.001). All pairs of genotyped individuals had negative or very low values of relatedness (ranging from 0.01 to -3.76), which excludes the possibility that we had sampled close relatives (Manichaikul et al. 2010).

**Table 1.**
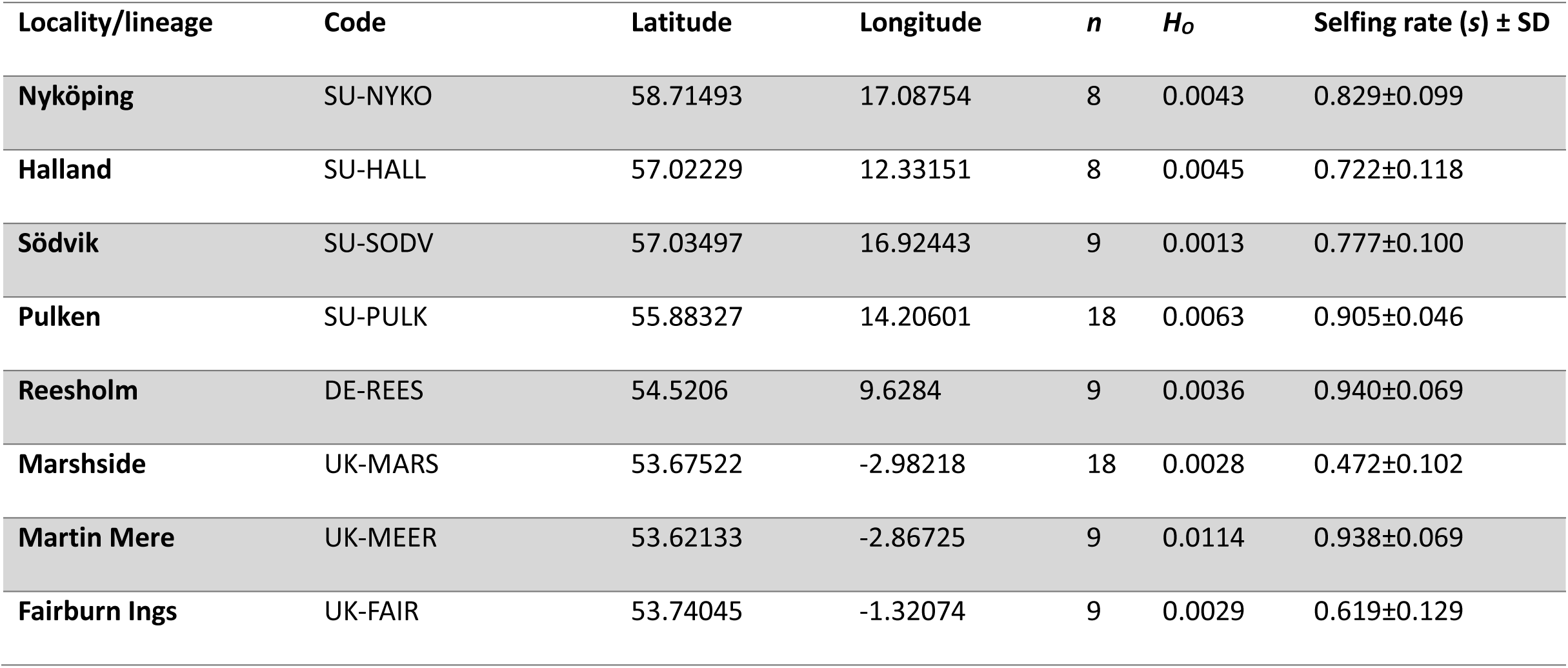
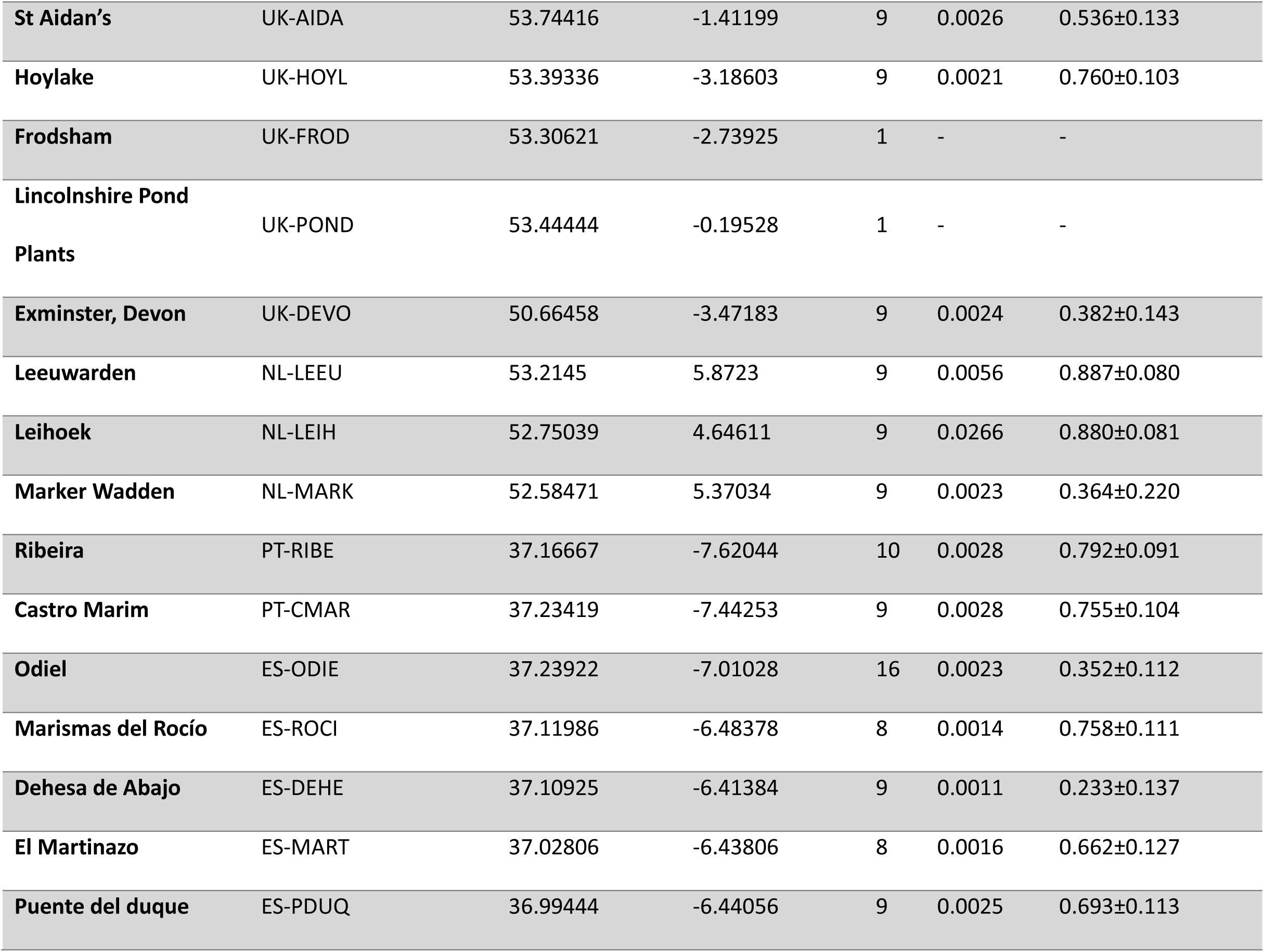
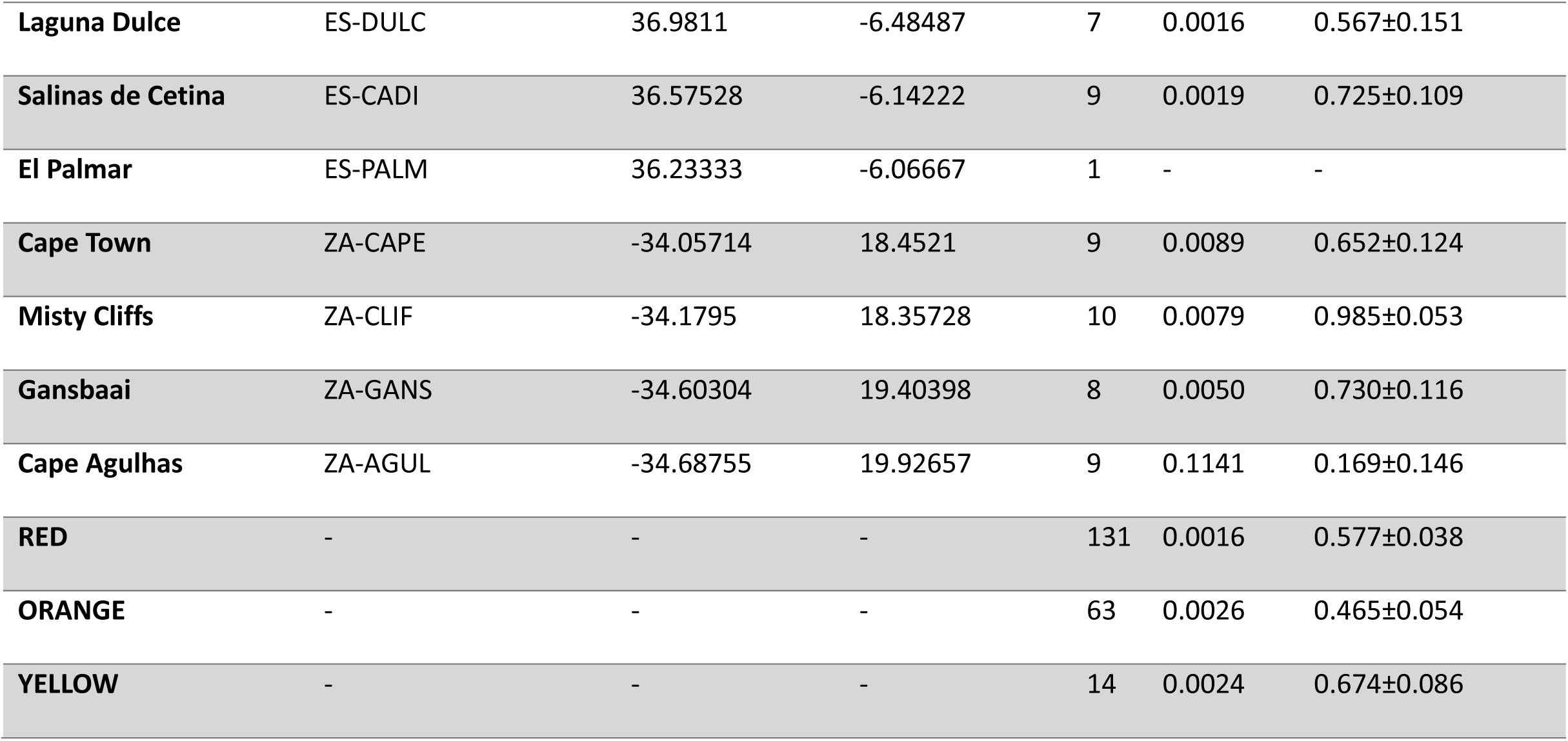
Locality/lineage, code, latitude, longitude, number of genotyped individuals (*n*), observed heterozygosity, self-fertilization rate (*s*), and its standard deviation (SD) for each sampled population and lineage (RED, ORANGE, and YELLOW) within the alien range as inferred by the Bayesian clustering method implemented in STRUCTURE. The first two characters of each locality indicate the corresponding country, following the International Organization for Standardization (ISO). Summary statistics for each lineage within the alien range were calculated only including non-admixed individuals (STRUCTURE *q*-values > 0.99).

### Population genetic structure and admixture

STRUCTURE v. 2.3.3 (Pritchard et al. 2000) analyses including all genotyped individuals from both Europe and South Africa (6,661 SNPs) identified *K*=2 as the most likely number of genetic clusters based on the Δ*K* criterion, while LnPr(X|*K*) reached a plateau at *K*=5 (Fig. S1a). For *K*>5, no further genetic structure was detected, yielding “ghost clusters” (i.e., clusters with no population or individual assigned to them; Guillot et al. 2005). For *K*=2, European individuals split into two main groups with low levels of admixture, whereas all South African populations are admixed between these two genetic clusters (Fig. S2). STRUCTURE analyses for *K*=5 showed the presence of three genetic clusters in Europe with limited correspondence with the geographical location of populations (Fig. 1b). One cluster (hereafter, RED lineage) was represented in multiple populations from Sweden, Germany, Netherlands, Spain, Portugal and one population from the UK, another cluster (hereafter, ORANGE lineage) was predominantly present in the UK and single populations from each of the Netherlands and Spain, and a third cluster (hereafter, YELLOW lineage) was represented in one population from each of Sweden and the Netherlands and some admixture in population PT-CMAR from Portugal (Fig. 2b). South African populations consisted of two genetic clusters, one of them represented in the population ZA-AGUL (hereafter, BLUE lineage) and another represented in the populations ZA-CLIF, ZA-GANS and ZA-CAPE. Some individuals scattered across different European populations showed a certain degree of admixed ancestry between the RED and YELLOW lineages or between the RED lineage and the BLUE lineage mostly present in the South African population ZA-AGUL. Only population PT-CMAR from Portugal showed a considerable degree of admixed ancestry between two lineages (RED × YELLOW) across most individuals. The commercial individual—acquired in the Lincolnshire Pond Plants Ltd. (Brookenby, UK)—from UK-POND was the only specimen showing evidence of admixture among three genetic clusters (ORANGE, YELLOW, and BLUE), suggesting a distinct genetic background compared to European field-sampled populations, likely associated with its horticultural origin. Among South African populations, only ZA-CAPE had an admixed ancestry between one lineage only present in South Africa and another (ORANGE) mostly present in European populations.

**Fig. 2.**
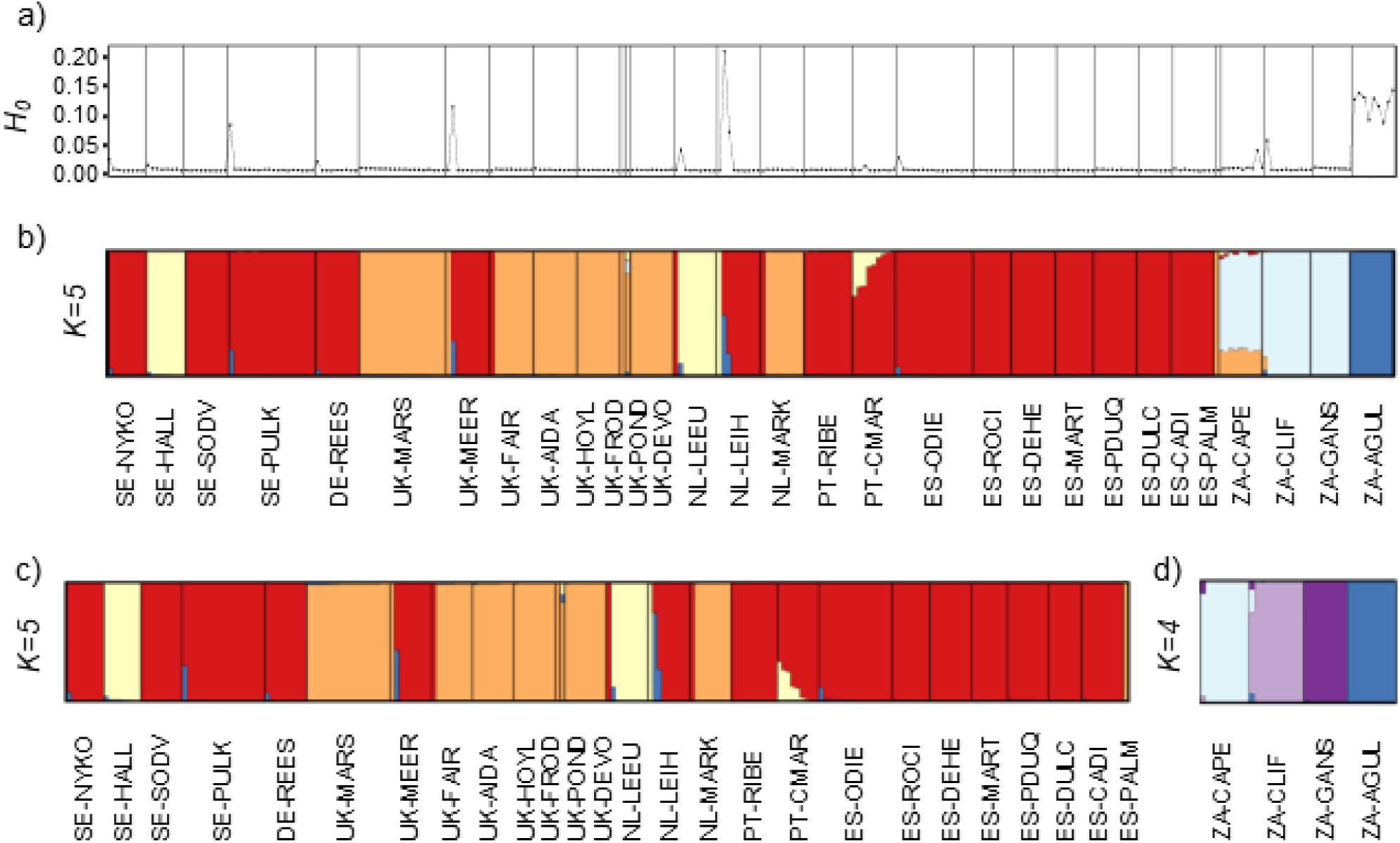
(a) Observed heterozygosity (*H_O_*) calculated for each individual of *C. coronopifolia*. Panels (b-d) show the results of genetic assignments based on the Bayesian clustering analyses implemented in STRUCTURE for datasets containing (b) all 266 individuals (6,661 SNPs), (c) 230 individuals from Europe (5,692 SNPs), and (d) 36 individuals from South Africa (13,079 SNPs). Each individual is represented by a vertical bar, which is partitioned into *K* coloured segments showing the individual’s probability of belonging to the cluster with that colour. Thin vertical black lines separate individuals from different populations. Population codes as described in Table 1.

STRUCTURE analyses exclusively focused on European populations (5,692 SNPs) identified *K*=2 as the most likely number of genetic clusters according to the Δ*K* criterion, whereas LnPr(X|*K*) reached a plateau at *K*=4 (Fig. S1b). Clustering solutions for *K*=2-4 are virtually identical to inferences obtained for STRUCTURE analyses including all genotyped individuals (see Fig. 2b-c, Fig. S3). STRUCTURE analyses focused on South African populations (13,079 SNPs) identified *K*=4 as the most likely number of genetic clusters according to the Δ*K* criterion, and LnPr(X|*K*) reached a plateau at the same *K* value (Fig. S1c). For *K*=2, we can see that the main separation is between ZA-AGUL and the other three populations (Fig. S4). For *K*=4, each population was assigned to a different genetic cluster, with very limited admixture in one individual from ZA-CAPE and another from ZA-CLIF (Fig. 2d).

### Phylogenomic analyses

Phylogenomic reconstructions in SVDQUARTETS (Chifman and Kubatko 2014) and dating analyses in BPP v. 4.1 (Flouri et al. 2018) suggested that the South African population ZA-AGUL diverged from a clade comprising all other lineages during the last glacial period (ca. 40 ka BP). This clade diverged almost simultaneously into five lineages around the Last Glacial Maximum (ca. 23 ka), which led to the formation of the rest of the South African lineages (ZA-GANS, ZA-CLIFF, and ZA-CAPE) and two European lineages, one of which split into the RED and YELLOW lineages during the Holocene (ca. 6 ka BP) (Fig. 3).

**Fig. 3.**
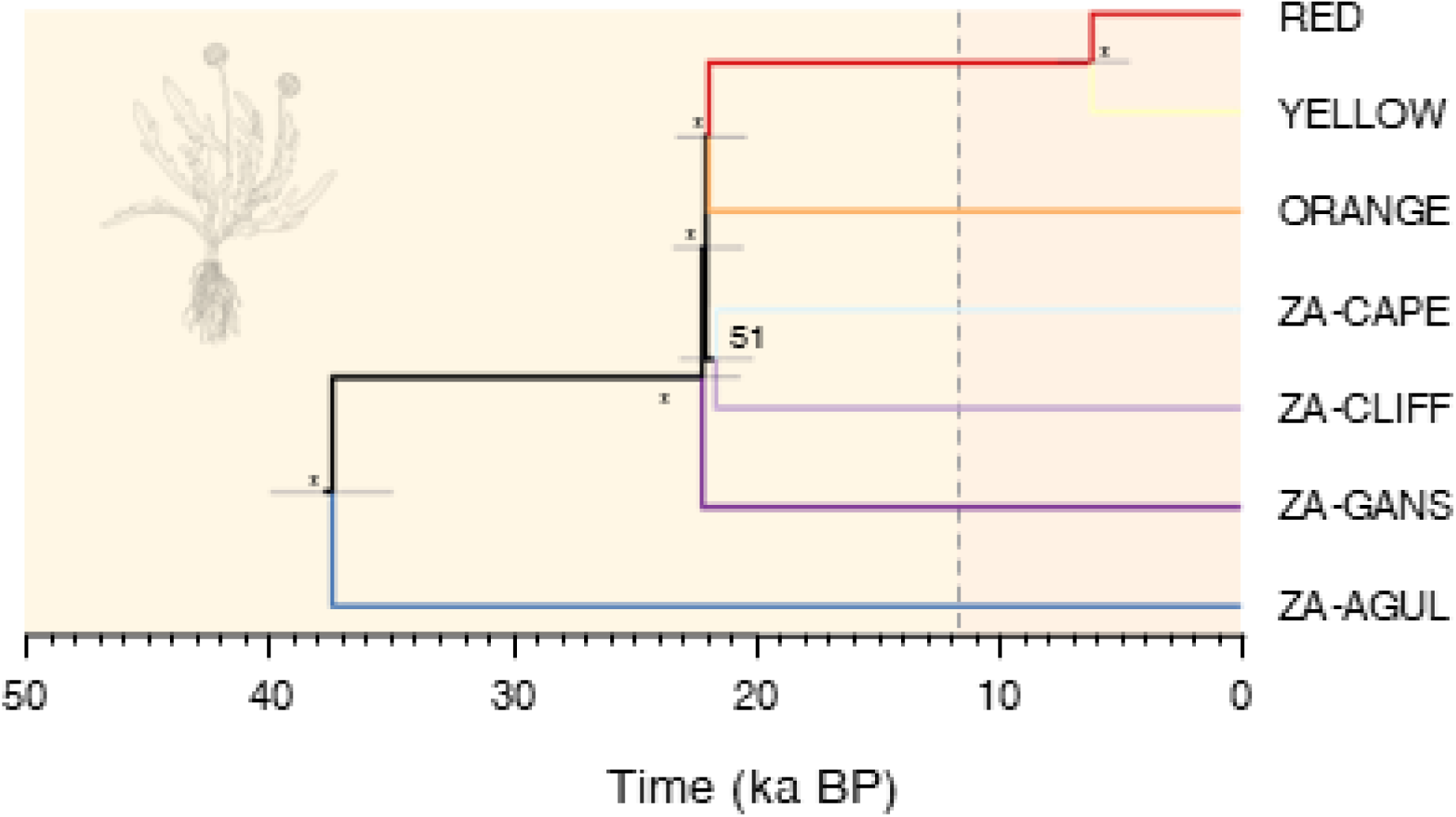
Phylogenetic tree inferred with SVDQUARTETS and divergence times estimated using BPP (analysis A00) for lineages of *C. coronopifolia* identified by the Bayesian clustering analyses implemented in STRUCTURE in the European invasive range (lineages RED, ORANGE and YELLOW) and the South African native range (populations ZA-CAPE, ZA-CLIFF, ZA-GANS, and ZA-AGUL). Bars on nodes indicate 95% highest posterior densities (HPD) of divergence times estimated considering a genomic mutation rate of 7.0 × 10^−9^ per site per generation and a one-year generation time; numbers on nodes indicate bootstrap support values (* = 100). Each colour matches with the inferred genetic clusters inferred by STRUCTURE analyses (see Fig. 2). Background colours indicate geological divisions of the Quaternary and the vertical dashed line separates the late Pleistocene (left) from the Holocene (right). Illustration of *C. coronopifolia*: Marina Trillo.

### Demographic history

STAIRWAY PLOT v. 2.1 analyses suggested that the three European lineages have experienced parallel demographic trajectories, with an abrupt demographic expansion preceding the documented arrival of the species to the continent, followed by demographic stagnation until the present day (Fig. 4a-c). Conversely, STAIRWAY PLOT suggests that most South African populations have experienced idiosyncratic demographic trajectories (Fig. 4d-g). The population ZA-CAPE went through a marked genetic bottleneck approximately 2 ka BP, which reduced its effective population size (*N_e_*) by > 90% with respect to the pre-bottleneck situation (Fig. 4d). The populations ZA-CLIFF and ZA-GANS experienced a sudden increase of *N_e_* during the Holocene (ca. 8 ka and 2 ka BP, respectively), followed by demographic stability (Fig. 4e-f). Finally, ZA-AGUL experienced a continuous decrease in *N_e_* since the onset of the Holocene until the present day (Fig. 4g).

**Fig. 4.**
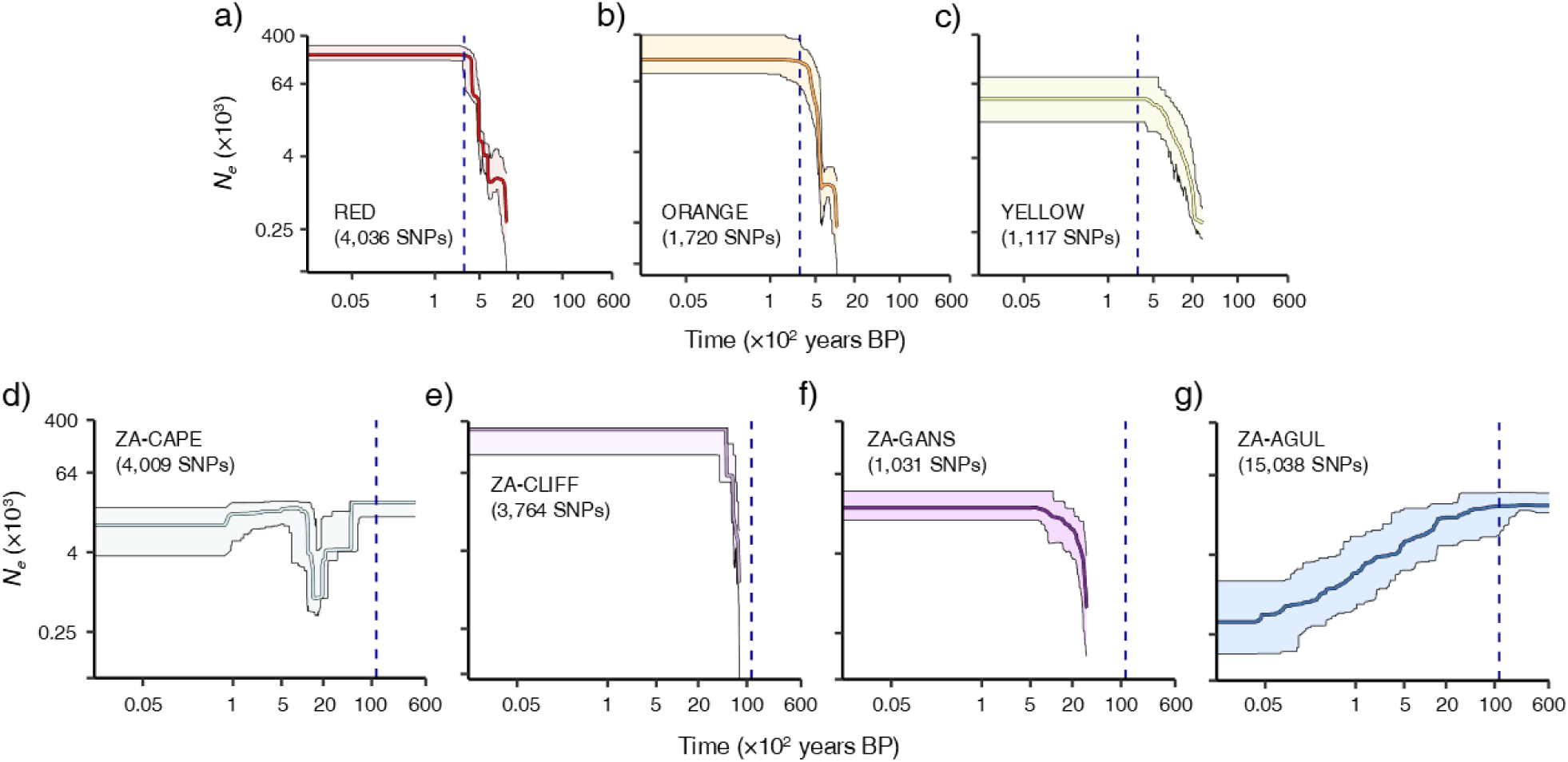
Demographic reconstructions inferred using STAIRWAY PLOT for each lineage of *C. coronopifolia* identified by Bayesian clustering analyses implemented in STRUCTURE for the European invasive range (lineages RED, ORANGE and YELLOW) and the South African native range (populations ZA-CAPE, ZA-CLIFF, ZA-GANS, and ZA-AGUL) (see Fig. 2). Plots show median (solid lines) and 2.5 and 97.5 percentiles (shaded areas) of effective population size (*N_e_*) through time, estimated assuming a genomic mutation rate of 7.0 × 10^−9^ per site per generation and one generation per year (both axes in logarithmic scale). Vertical dashed lines indicate the first record of the presence of the species in Europe (1739, 286 years ago) for the European lineages and the onset of the Holocene (ca. 11,650 BP) for the South African populations.

### Estimates of population genetic diversity and selfing rates

Genetic diversity was very low across most genotyped individuals and populations (Table 1 and Fig. 2). The only exceptions were some admixed individuals and the native ZA-AGUL population, which exhibited markedly higher levels of genetic diversity (Table 1 and Fig. 2a). When excluding these, genetic diversity —estimated as observed heterozygosity (*H_O_*)— was significantly higher in populations from the native range than in those from the invasive range in Europe (Wilcoxon rank-sum test, *W* = 651; *n* = 208, 26; *p* < 0.001).

When admixed individuals were excluded, levels of genetic diversity differed significantly among the three main lineages identified in the invasive range (Kruskal–Wallis rank sum test, *χ²* = 55.43; *df* = 2; *n* = 131, 63, 14; *p* < 0.001). Post hoc pairwise Wilcoxon rank-sum tests indicated that individuals assigned to lineage ORANGE had significantly higher genetic diversity than those in lineage RED (*p* < 0.001) but did not differ significantly from those in lineage YELLOW (*p* = 0.402). Genetic diversity did not differ significantly between the RED and YELLOW lineages (*p* = 0.083).

Admixed individuals, especially those involving the RED and BLUE lineages, had higher levels of genetic diversity compared to non-admixed individuals assigned to either of these two genetic clusters (Wilcoxon rank-sum test, *W* = 949; *n_1_* = 134, *n_2_* = 9; *p* < 0.001). However, individuals from population PT-CMAR, which exhibits considerable genetic admixture between the RED and YELLOW lineages, did not show higher levels of genetic diversity compared to non-admixed European individuals assigned to the same genetic clusters (Wilcoxon rank-sum test, W = 772; *n* = 131, 14; *p* = 0.426).

Selfing rates (s) were significantly distinct from random for most sampled populations (*H_0_* = s = 0; *p* < 0.001), with the exception of NL-MARK (*p* = 0.221) and ZA-AGUL (*p* = 0.149) (Table 1). Selfing rates ranged from 23.3% in ES-DEHE to 98.5% in ZA-CLIF, indicating a high prevalence of selfing across most populations (Table 1). RMES analyses conducted by pooling all non-admixed (STRUCTURE *q*-values > 0.999) individuals within each lineage inferred by STRUCTURE in Europe also revealed high selfing rates (RED: 57.7%; ORANGE: 46.5%; YELLOW: 67.4%; Table 1).

### Hybrid simulations for the determination of recency of admixture between lineages

Hybrid simulations between lineages RED and YELLOW indicate that only one individual from PT-CMAR had levels of genetic diversity and admixture compatible with a backcross between F1 and the parental RED lineage (Fig. 5b). The heterozygosity for the rest of individuals from PT-CMAR was much lower than expected for any of the simulated hybrid classes with comparable levels of admixed ancestry (Fig. 5b). Hybrid simulations between lineages RED and BLUE demonstrated that most genotyped individuals exhibited levels of genetic diversity compatible with those obtained for simulated hybrid classes with similar admixed ancestries (Fig. 5c).

**Fig. 5.**
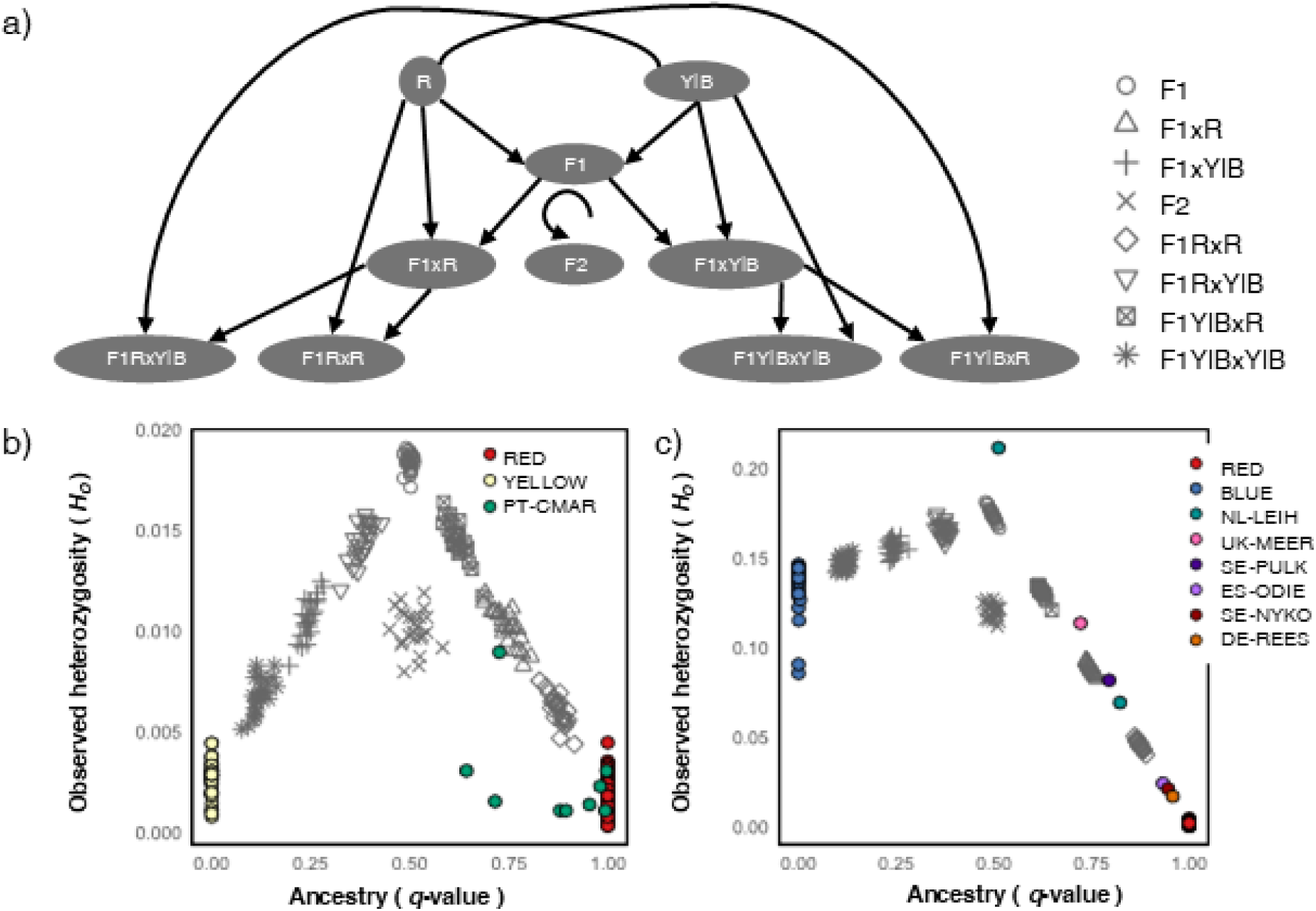
Simulation of different hybrid classes for crosses between lineages. Panel (a) shows a schematic of simulated crosses (R=RED lineage, Y = YELLOW, and B= BLUE). Panels (b) and (c) show observed heterozygosity (*H_O_*) plotted against ancestry inferred by STRUCTURE analyses for observed (coloured dots) and simulated (grey shapes) genotypes in crosses between (b) lineages RED and YELLOW and (c) lineages RED and BLUE. Panels (b) and (c) show pure individuals for each lineage (RED, YELLOW, and BLUE) in our empirical dataset, whereas sampled individuals presenting an admixed ancestry are assigned to their respective populations using different colour codes.

## DISCUSSION

By using *C. coronopifolia* as a model system to test Baker’s “ideal weed” hypothesis, our study provides one of the first population-genomic analysis of the colonization history and successful spread of a highly invasive selfing species. Our results reveal that the species’ invasion success in Europe is probably linked to a demographic scenario shaped by strong selfing. The predominantly self-fertilizing mating system, previously suggested but never quantified, has contributed to generate extremely low within-population diversity and pronounced genetic differentiation across both native and introduced ranges. These findings are consistent with the predictions of the “ideal weed” framework, which postulates that reproductive assurance through selfing can enhance colonization ability by enabling single individuals to establish viable populations. At the same time, the genetic structure observed among European populations reflects introductions of multiple lineages followed by rapid expansion under selfing, illustrating how reproductive assurance can compensate for low propagule diversity and facilitate large-scale range expansion. Altogether, these results position selfing as a major driver of the genetic landscape of *C. coronopifolia*, which explains observed patterns of genetic diversity and structure and provides important clues to understand the colonization history and successful spread.

### Genetic structure and the origin of invasive lineages

Bayesian clustering analyses revealed a marked genetic structure in both the native and invasive ranges, likely resulting from high selfing rates and extremely limited gene flow among populations and across lineages (Fig. 2). Each of the genotyped populations from the native range of *C. coronopifolia* in South Africa formed a distinct genetic cluster, with very limited signatures of admixture even among populations separated by short geographical distances (e.g., <20 km between ZA-CAPE and ZA-CLIFF; Fig. 1). In the invasive European range, we identified three genetic clusters distributed across different regions of the continent. With a few exceptions, most populations were dominated by individuals belonging to the same genetic cluster. This pattern may reflect a priority effect, whereby first-established lineages reduce the chances of later arrivals to successfully establish, reinforcing the persistence of genetically homogeneous populations and limiting admixture (Fraser et al. 2015).

None of the three lineages found in Europe was detected in South Africa, suggesting that they either originated in the invasive range (e.g., via genetic drift fuelled by selfing) or, more likely, derive from the source populations that had remained uncovered by our sampling in the native range. Phylogenomic reconstructions and divergence time estimates provide important insights into the origin of the European lineages. Dating analyses indicate that ZA-AGUL diverged from other lineages approximately 40 ka BP, while the remaining South African populations and European lineages split around 22 ka BP, with the most recent split, between the RED and YELLOW European lineages, taking place ca. 6 ka BP. These divergence times (>6 ka) largely predate the first European records of the species in the 18th century, indicating that the three lineages in the invasive range did not originate through in situ diversification and, instead, most likely colonized Europe after one or more introduction events from unsampled source populations in the native range. The pronounced genetic structure of the species highlights the challenge of identifying the specific ancestral sources of European lineages, a task that would likely require extensive fine-scale sampling across the native range.

### Demographic history

Demographic reconstructions in STAIRWAY PLOT show that the four South African populations experienced independent demographic trajectories, which is not surprising given their marked genetic differentiation and distinct reproductive systems (i.e., demographic independence). Although the European lineages were not detected in our limited sampling from South Africa, and direct comparisons of genetic diversity between native and invasive populations are therefore difficult, non-admixed individuals from Europe showed significantly lower levels of heterozygosity than non-admixed individuals from the predominantly selfing populations in the native range (i.e., ZA-CAPE, ZA-CLIF, and ZA-GANS), suggesting that the colonization of Europe further contributed to the loss of genetic diversity in the invasive populations. However, given that the source populations were likely already genetically depauperate, the extent to which the colonization process further reduced genetic diversity in European populations appears to be moderate (38.4% relative to native selfing populations), corresponding to a decrease in mean observed heterozygosity (*H_O_*) from 0.005 to 0.002. Accordingly, all three European lineages exhibited a sudden increase in effective population size (*N_e_*) beginning around 300 years ago, roughly coinciding with their introduction to Europe, followed by demographic stagnation up to the present day. Because very low genetic diversity in highly selfing species can distort coalescent patterns and may violate the assumptions of demographic models, such reconstructions should be interpreted with caution (Hartfield 2017). Nevertheless, our results suggest that selfing may have minimized founder effects during introduction and led to a constant effective population size (*Ne*) during the invasion, despite census population sizes (*Nc*) expanded rapidly in newly colonized areas. This pattern contrasts with expectations for highly invasive outcrossing organisms, which typically experience a genetic bottleneck after introduction followed by explosive demographic expansions during the invasion phase (e.g., Ortego et al. 2021).

### Introduction and invasion history

The three lineages recovered in the invasive range of *C. coronopifolia* in Europe are distributed across different regions in the continent, with limited congruence between genetic structure and geography. This pattern likely reflects the interplay between introduction history and subsequent expansion facilitated by long-distance dispersal (LDD). The earliest records of *C. coronopifolia* in Europe (Germany: 1739; Netherlands: 1742) suggest an initial introduction via ship ballast water after the Dutch colonized South Africa in 1652. Recently, Verloove and Gonggrijp (2025) speculated that this plant may have colonized Europe naturally from Africa, with a south-to-north colonization pathway (e.g., from the Iberian Peninsula to Scandinavia), in which case we would expect to find all lineages in southern Europe and only a subset in more recently colonized northern regions. However, our data do not support this scenario and, instead, point to multiple introductions and the arrival of at least three different lineages, all currently present in the Netherlands, from where they may have spread to the rest of Europe.

The expansion of the species may also have been reinforced by its ongoing sale as an ornamental pond plant, with a reported deliberate introduction into gardens in the vicinity of UK-HOYL in the late 19^th^ century (Wilkinson 2023). However, we found that the commercial plant that we genotyped (UK-POND) showed an admixed ancestry involving three lineages, in contrast with field-sampled individuals, which were either fully assigned to a single genetic cluster or exhibited admixture between two of them. This suggests that horticulture has contributed to admixture in the commercially available plants that is not represented in any of the field sites we have sampled, suggesting that deliberate spreading of ornamental plants has probably not played an important role in the European expansion of this alien species. In contrast, there is good evidence for the effective dispersal of *C. coronopifolia* by gut passage inside migratory waterbirds (i.e., endozoochory) into wetlands with a broad salinity range (Raulings et al. 2011; Lovas-Kiss et al. 2019; Sánchez-García et al. 2024). Previous studies have found evidence that migratory waterbirds shape genetic connectivity for native plants in Europe (Green et al. 2023), and it is likely that avian vectors have also played a key role in the invasion success of *C. coronopifolia*. The Netherlands, where the three European lineages are present, acts as a hub for many migratory bird species providing strong connections with all the other European areas (Delany et al. 2009). Accordingly, most of the field sites we sampled hold major concentrations of migratory birds and are protected for that reason. Seed dispersal by floating along currents could also have an important role in colonization along coastlines (Costa et al. 2009), whereas livestock may also carry seeds both externally and through gut passage (Tomasson 2020).

### A predominantly selfing mating system

Our results reveal a predominantly selfing mating system in *C. coronopifolia*, with a mean self-fertilization rate of ∼70% (ranging from 23.3% to 98.5%; Table 1). As frequently found in other flowering plants, our results indicate predominant but not obligate selfing, with the presence of variable rates of outcrossing across populations (Siol et al. 2008; Whitehead et al. 2018; Jullien et al. 2021). One remarkable exception is the native population ZA-AGUL from South Africa, where selfing was undetectable and which presented much higher levels of genetic diversity compared to the rest of the populations (Table 1; Fig. 1a). Phylogenomic reconstructions and dating analyses showed that the lineage present in this population was the earliest diverged (∼40 ka BP; Fig. 4), suggesting an evolutionary transition to selfing from an ancestral outcrossing or predominantly outcrossing mating system (Ortiz et al. 2007). Accordingly, the evolution of selfing from obligate cross-fertilization is the most frequent reproductive transition in angiosperms and has occurred repeatedly in different lineages over a range of contrasting evolutionary timescales (Stebbins 1957; Barrett 2008; Cutter 2019; e.g., genus *Primula*: Wang et al. 2021; genus *Capsella*: Slotte et al. 2012; Brandvain et al. 2013). In the same vein, the lack of selfing in the European population NL-MARK, belonging to a lineage in which the rest of the populations present high selfing rates, indicates that the reproductive mode is probably highly labile and context dependent (Whitehead et al. 2018). Thus, our results suggest the presence of different reproductive strategies across and within lineages and populations of *C. coronopifolia*, in line with plasticity in mating-system documented in other flowering plants (Whitehead et al. 2018; Suijkerbuijk et al. 2025; e.g., *Arabidopsis lyrate*: Mable and Adam, 2007; Willi and Määttänen, 2010). Multiple factors could explain contrasting selfing rates across populations and local selection for a particular reproductive strategy, including relative abundances of efficient pollinators or different sources of abiotic, biotic or human-induced stressors, among others (Mable and Adam, 2007; Willi and Määttänen 2010; Whitehead et al. 2018; Cutter 2019; Suijkerbuijk et al. 2025). In addition, populations with high-selfing rates are more likely to undergo effective dispersal by birds and colonize new sites via a single seed, which may represent an important evolutionary advantage during range expansions following initial colonization of non-native areas (Baker 1955; Razanajatovo et al. 2019).

The consequences of selfing are evident in observed patterns of genetic diversity and structure across populations of *C. coronopifolia*. Selfing is well known to reduce within population genetic diversity while amplifying genetic differentiation among populations, effects that are patent in our data and mirror previous studies where selfers often show much lower heterozygosity and higher structure relative to outcrossers (Charlesworth and Wright 2001; Glémin et al. 2006; Siol et al. 2008; Glémin and Ronfort 2013; Hartfield et al. 2017). Accordingly, we found consistently very low levels of genetic diversity across all selfing populations, which contrast with the higher diversity in population ZA-AGUL, where outcrossing dominates and selfing is probably absent or very limited (Table 1; Fig. 1a). Genetic diversity in population NL-MARK, where we found no significant selfing rates, was comparable to that of predominantly selfing populations. This suggests that outbreeding had no positive impact on the genetic diversity of this population, likely because it originated from an already genetically depauperate selfing population and has experienced no gene flow from other populations with different genetic backgrounds.

Selfing also appears to be the main driver of observed genetic structure in *C. coronopifolia*, with three and four distinct lineages in Europe and South Africa, respectively, each largely isolated and with limited admixture. This suggests that selfing has facilitated lineage diversification by limiting gene flow and maintaining distinct genetic clusters over short geographic scales, even allowing their co-existence within certain populations (Loveless and Hamrick 1989; Wright et al. 2013; Hu 2015). Accordingly, our data revealed the co-occurrence of different lineages in several invasive populations in Europe (UK-MEER, UK-FAIR, NL-LEUU, NL-LEIH, and NL-MARK; Fig. 1), where individuals assigned to different genetic clusters co-existed without evidence of genetic admixture (Fig. 2).

Although selfing seems to dominate in most populations, our Bayesian clustering analyses also revealed the presence of some admixed individuals resulted from sporadic outcrossing events involving different lineages (Fig. 2b). As expected, many of these individuals presented substantially higher levels of heterozygosity compared to non-admixed individuals (Fig. 2a), illustrating how outcrossing events can rapidly increase individual genetic diversity. However, our data also showed that several individuals with admixed genetic ancestry exhibit heterozygosity levels as low as those of non-admixed individuals from the parental selfing lineages. Simulations further indicated that, although initial admixture can elevate genetic diversity, successive generations of selfing rapidly reduce heterozygosity, eventually restoring values comparable to non-admixed selfing populations. This is the case of PT-CMAR population, which includes individuals with varying levels of admixture between RED and YELLOW lineages (Fig. 2c), yet shows levels of heterozygosity as low as those found in non-admixed populations from either parental lineage (Fig. 5b). Although the increased genetic diversity in first generation admixed individuals is often diminished by subsequent selfing, our data also demonstrate effective allele exchange across selfing lineages, potentially enabling adaptive gene flow if introgressed variants provide local fitness advantages (Suarez-Gonzalez et al. 2018; Burgarella et al. 2019; Lewis et al. 2023).

## CONCLUSIONS

In summary, our population-genomic analyses reveal that the invasion success of *C. coronopifolia* in Europe is probably linked to its colonization history and the predominance of selfing as a reproductive strategy. The species’ genetic makeup —characterized by a strong population structure and limited gene flow— suggests multiple introductions, or at least the introduction of multiple lineages followed by rapid expansion under a selfing regime. This combination of reproductive assurance and dispersal versatility has enabled *C. coronopifolia* to establish and persist across diverse European habitats despite its reduced genetic diversity. By placing these findings within Baker’s framework of the “ideal weed” and the “general-purpose genotype”, our study highlights how traits such as self-compatibility, vegetative reproduction, and ecological generalism can overcome the genetic constraints typically associated with colonization bottlenecks. Our study exemplifies how mating system evolution can mediate the balance between colonization ability and long-term adaptability during invasion. Future research should expand this framework by incorporating whole-genome sequencing across native and invaded ranges to determine the genomic basis of alternative mating systems, assess adaptive differentiation among lineages, and explore the evolutionary consequences of selfing, including the purging of deleterious alleles and the development of selfing-syndrome traits (Barrett 2008; Brandvain et al. 2013; Hartfield 2016). Comparing European populations with those from other invaded continents will help disentangle primary versus secondary colonization pathways, whereas investigating the role of migratory waterbirds and livestock as potential dispersal vectors will further clarify how the interplay between natural and anthropogenic processes have shaped the species’ global spread.

## MATERIALS AND METHODS

### Plant material and sampling sites

From July 2021 to April 2023, we sampled thirty populations of *C. coronopifolia* across its native and alien range in South Africa (*n* = 4) and Europe (*n* = 26), respectively (Table 1). We collected mature plants along transects with a spacing of 1-3 metres between individuals and preserved them in silica gel until needed for DNA extraction. Additionally —to see if there are any differences between plants collected in the wild and those from horticulture— we acquired a cultivated individual from the Lincolnshire Pond Plants Ltd. (Brookenby, UK). Five individuals of *Cotula* spp. collected in Knysna, Leisure Island (-34.06517, 23.06142), South Africa, were used as an outgroup in phylogenomic analyses. We used a global positioning system (GPS) or Google Earth to record spatial coordinates for each site. Geographical coordinates and other details of sampling sites are indicated in Table 1.

### Sequencing and processing of genomic data

We used the DNeasy Plant Mini Kit (Qiagen, Valencia, CA, USA) to extract and purify DNA from leaf tissue, according to the manufacturer’s instructions. We determined genomic integrity on 1% agarose gels and quantified DNA concentration using a Qubit Fluorometer (Thermo Fisher Scientific®). Genotyping by sequencing (GBS) libraries were prepared by Ecogenics GmbH (Switzerland) according to Elshire et al. (2011) and using the restriction enzymes *EcoRI* and *MseI*. The resulting PCR products were sequenced on an Illumina NovaSeq platform using an S2 flow cell (2×100 bp).

Demultiplexed sequence data were assembled de novo using default parameters in IPYRAD v. 0.9.93 (Eaton and Overcast, 2020). The filtering step was performed by allowing a maximum of five low-quality base calls per read and a minimum read length of 35 bp after adapter trimming. For alignment, we used a clustering threshold of 90%, allowing a maximum of two alleles per site in the consensus sequences. Unless otherwise indicated, all downstream analyses were performed using unlinked (one SNP per locus) single nucleotide polymorphism (SNP) data sets. We used the option *relatedness2* in VCFTOOLS v. 0.1.17 to calculate the relatedness among all pairs of genotyped individuals and to exclude the possibility that we had sampled close relatives within each study population (Manichaikul et al. 2010; Danecek et al. 2011).

### Population genetic structure and admixture

We assessed population genetic structure and admixture using the Bayesian Markov chain Monte Carlo clustering method implemented in the program STRUCTURE v. 2.3.3 (Pritchard et al. 2000). To fully explore population genetic structure and admixture, we initially analysed the data from all populations jointly and, subsequently, we ran independent analyses exclusively focused on populations from South Africa (native range) and Europe (invasive range). We ran STRUCTURE analyses assuming correlated allele frequencies and admixture and without using prior population information (Hubisz et al. 2009). To estimate the most likely number of genetic clusters, we ran several independent runs for each value of *K* (from *K* = 1 to *K* = 8) with 200,000 MCMC cycles, following a burn-in step of 100,000 iterations. The 10 runs with the highest likelihood for each value of *K* were retained and determined the number of genetic clusters that best described our data according to the log probabilities of the data (LnPr(X|*K*) (Pritchard et al. 2000) and the Δ*K* method (Evanno et al. 2005), as implemented in STRUCTURE HARVESTER (Earl and vonHoldt 2012). We used PONG v. 1.5 (Behr et al. 2016) to visualize the cluster membership of individuals as bar plots.

### Phylogenomic analyses

First, we used matrices of unlinked SNPs to reconstruct the phylogenomic relationships among lineages in SVDQUARTETS (Chifman and Kubatko 2014). We included five representative individuals for each lineage identified by STRUCTURE analyses (STRUCTURE *q*-values > 0.999; see Results section), used five individuals of *Cotula* spp. as an outgroup, exhaustively evaluated all possible quartets, and performed nonparametric bootstrapping with 100 replicates to quantify uncertainty in relationships. Second, we used the multispecies coalescent model implemented in the program BPP v. 4.1 (Flouri et al. 2018) to estimate divergence times (analysis A00). The *.loci* file from IPYRAD was edited and converted into a BPP input file using custom scripts (J. Ortego; available from https://github.com/OrtegoLab/ipyrad2bpp). Due to high computational demands, we only considered a subset of 1,000 loci for BPP analyses. We set the phylogenetic tree inferred with SVDQUARTETS as tree prior, applied automatic adjustment of fine-tune parameters, set the diploid option to indicate that the input sequences are unphased, and adjusted the inverse-gamma distributions of the θ (α = 3, β = 0.04) and *τ* (α = 3, β = 0.07) priors according to empirical estimates calculated from the number of segregating sites per site (Huang et al. 2020). We ran two independent replicates for 1,000,000 generations, with samples taken every 2 generations, after a burn-in of 100,000 generations. The estimation of divergence times was conducted using the equation τ = 2*μ*t, where τ indicates the divergence in substitutions per site as estimated by BPP, *μ* refers to the per-site mutation rate per generation, and t represents the absolute divergence time in years (Huang et al. 2020). We considered the mutation rate per site per generation of 7.0 × 10^−9^ estimated for *Arabidopsis thaliana* (Ossowski et al. 2010).

### Demographic history

We reconstructed the past demographic history of each lineage using STAIRWAY PLOT v. 2.1, an approach which implements a flexible multi-epoch demographic model based on the site-frequency spectrum (SFS) and that does not require whole-genome sequence data or reference genome information (Liu and Fu 2020). We calculated the SFS for each lineage inferred by STRUCTURE (see Results and Fig. 2) and ran STAIRWAY PLOT considering one generation per year, the mutation rate of 7.0×10^-9^ per site per generation estimated for *Arabidopsis thaliana* (Ossowski et al. 2010), and 200 bootstrap replicates to estimate 95% confidence intervals.

### Estimates of population genetic diversity and selfing rates

To quantify the genetic diversity, we calculated observed heterozygosity (*H_O_*) for each individual as implemented in the R package *hierfstat* (Goudet 2005). Then, we ran Wilcoxon–Mann-Whitney tests in R to compare *H_O_* values between pure and admixed individuals identified by STRUCTURE analyses (see Results). Selfing rates were calculated for each population (see Table 1) and European lineages inferred by STRUCTURE analyses (STRUCTURE *q*-values > 0.999) using the robust multilocus estimate of selfing (RMES), with 10,000 iterations to generate the *p*-values (David et al. 2007; Miller and Coltman 2014). Since RMES software uses the heterozygosity variance across the target population, we generated a separate dataset for each population or lineage. Due to computational limitations, we selected 1,000 random SNPs for each dataset (e.g., Adhikari et al. 2021).

### Hybrid simulations to determine the recency of admixture between lineages

STRUCTURE analyses revealed that some individuals present different degrees of admixed ancestry, suggesting either contemporary or historical hybridization between lineages (see Results). More specifically, admixed individuals primarily showed different ancestry proportions involving either two lineages distributed in Europe or one lineage distributed in Europe and another distributed in South Africa (see Results). To evaluate whether admixed individuals match or deviate from genetic diversity and admixture expectations corresponding to different hybrid classes resulting from recent hybridization between lineages (i.e., F1, F2 and different backcrosses), as opposed to historical events, we performed hybrid simulations using HYBRIDLAB (Nielsen et al. 2006); e.g., Salces-Castellanos et al. 2020. We performed independent simulations considering the two pairs of lineages primarily involved in the formation of admixed individuals (RED and YELLOW; RED and BLUE). We selected as reference parental lineages pure individuals with ≥ 99.9% probability of cluster membership, as determined by the STRUCTURE analyses on our empirical dataset (see Results and Fig. 2) and used them to simulate 25 individuals corresponding to each of six different hybrid classes, as illustrated in Fig. 3a. Finally, we ran STRUCTURE analyses for datasets including both empirical and simulated individuals, calculated their admixture coefficients and plotted them against their respective estimates of observed heterozygosity (*H_O_*; calculated as detailed previously). If admixture is the consequence of recent hybridization between lineages, we would expect that patterns of genetic diversity and admixture of empirical individuals align to those expected for a particular simulated hybrid class. On the contrary, if observed admixture is the outcome of historical hybridization, we would expect lower levels of genetic diversity than expected from admixture proportions for a given simulated hybrid class, especially if several generations of selfing have taken place after the hybridization event.

## AKNOWLEDGEMENTS

This research was funded by Ministerio de Ciencia e Innovacion, project number PID2020–112774GB-I00/AEI/10.13039/501100011033, and RSG was funded by an FPI grant from the Spanish Ministerio de Ciencia, Innovacion y Universidades (PRE2021-099466). PT sampling was funded by FCT – Foundation for Science and Technology UID/04326/2025 and LA/P/0101/2020. Sampling permits were provided by the Junta de Andalucía (project 2021/13), English Nature, The Wildfowl & Wetlands Trust, The Royal Society for the Protection of Birds, Salinas de Cetina and Instituto da Conservacão da Natureza e Florestas, and all necessary sampling permissions were obtained prior to sampling elsewhere. Logistic support was provided by Laboratorio de Ecología Molecular (LEM-EBD) from Estación Biológica de Doñana (CSIC). We thank Prof. Glynis Goodman-Cron for the taxonomic clarification of the specimens collected in South Africa. We also thank Centro de Supercomputación de Galicia (CESGA) and Doñana’s Singular Scientific-Technical Infrastructure (ICTS-RBD) for access to computer resources.

## AUTHOR CONTRIBUTIONS

RSG, JO, MAO and AJG conceived and designed the study. RSG, AJG, MAO, FH, CR, JR, EAS, LT, KT and CHAL collected the samples. RSG analysed the data, with input from JO. RSG wrote the first draft of the manuscript. JO and AJG revised the first and later drafts. MAO, CG, EAS, KT and CHAL contributed to later drafts. All authors read and approved the final manuscript.

## Data Availability

Raw Illumina reads have been deposited at the NCBI Sequence Read Archive (SRA) under BioProject XXX. Input files for all analyses are available for download on Figshare (https://doi.org/XXX).

