## Supplementary Material for "Genomic footprints of selfing, introduction history, and long-distance dispersal in an invasive alien plant"

Raúl Sánchez-García[1*], Andy J. Green[1,2], María A. Ortiz[3], Cristina García[4], Francisco Hortas[5], Chevonne Reynolds[6], Jennifer Rowntree[7], Ester A. Serrão[8], Lina Tomasson[9], Karin Tremetsberger[10], Casper H.A. van Leeuwen[11], and Joaquín Ortego[12]

[1] Department of Conservation Biology and Global Change, Doñana Biological Station (EBD), CSIC, Seville, Spain

[2] Department of Natural Sciences, Manchester Metropolitan University, Manchester, UK

[3] Department of Plant Biology and Ecology, University of Sevilla, Seville, Spain

[4] Department of Biological Sciences, School of Life and Environmental Sciences, Royal Holloway University of London, Egham, Surrey, UK

[5] Department of Biology, Institute of Marine Research (INMAR), University of Cadiz, Puerto Real, Spain

[6] School of Animal, Plant and Environmental Sciences, University of the Witwatersrand, Johannesburg, South Africa

[7] School of Biological and Marine Sciences, University of Plymouth, Drake Circus, Plymouth, UK

[8] Centre of Marine Sciences, CCMAR, University of Algarve, Campus de Gambelas, Faro, Portugal

[9] National Coordinator for Aquatic Invasive Alien Species, Swedish Agency for Marine and Water Management, Gothenburg, Sweden

[10] BOKU University, Institute of Botany, Department of Ecosystem Management, Climate and Biodiversity, Vienna, Austria

[11] Department of Ecology, Radboud Institute for Biological and Environmental Sciences, Radboud University, Nijmegen, the Netherlands

[12] Department of Ecology and Evolution, Doñana Biological Station (EBD), CSIC, Seville, Spain

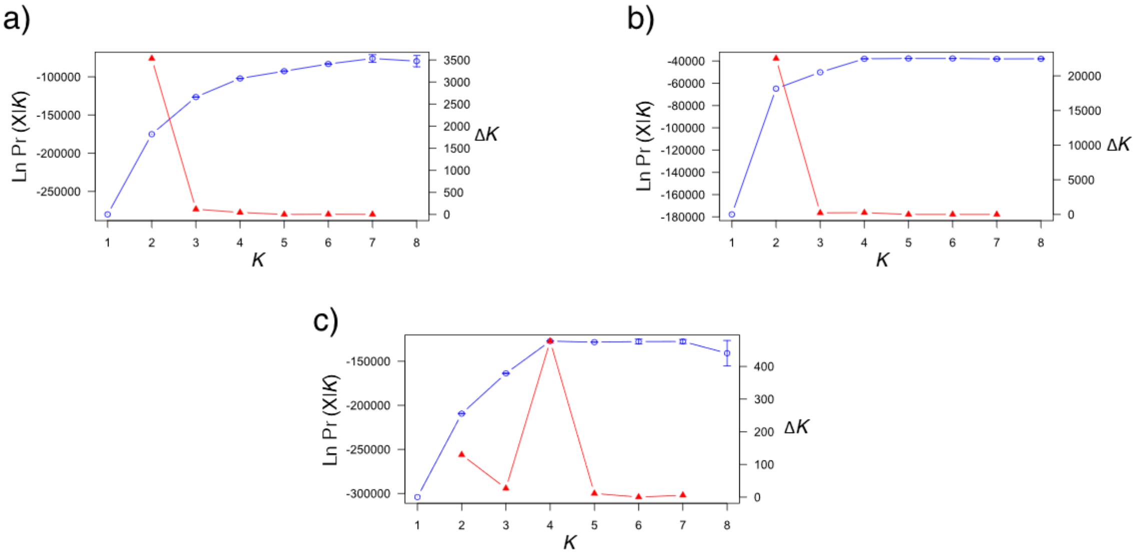


Fig. S1. Mean (±SD) log probability of the data (LnPr(X|*K*)) over 10 runs of STRUCTURE (left axes, blue dots and error bars) for each value of *K* and the magnitude of Δ*K* (right axes, red triangles) for (a) dataset containing all 266 individuals, (b) 230 individuals from Europe, and (c) 36 individuals from South Africa.

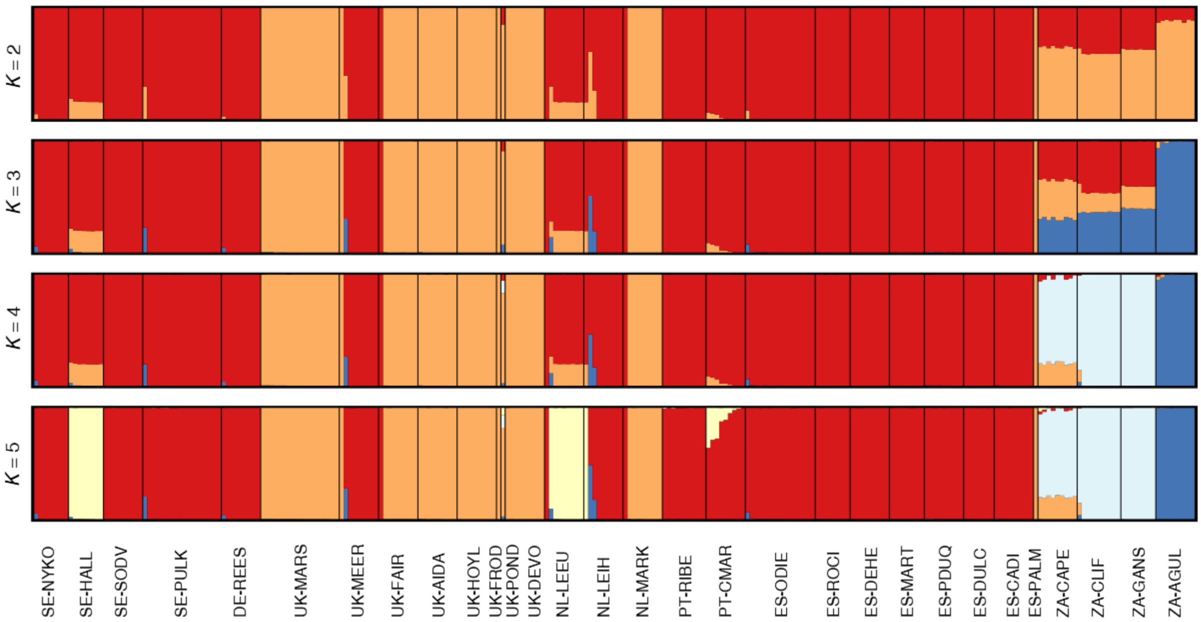


Fig. S2. Results of genetic assignments based on the Bayesian clustering analyses implemented in STRUCTURE (*K* = 2-5) for the dataset containing all 266 individuals (6,661 SNPs). Each individual is represented by a vertical bar, which is partitioned into *K* coloured segments showing the individual's probability of belonging to the cluster with that colour. Thin vertical black lines separate individuals from different populations. Population codes as described in Table 1.

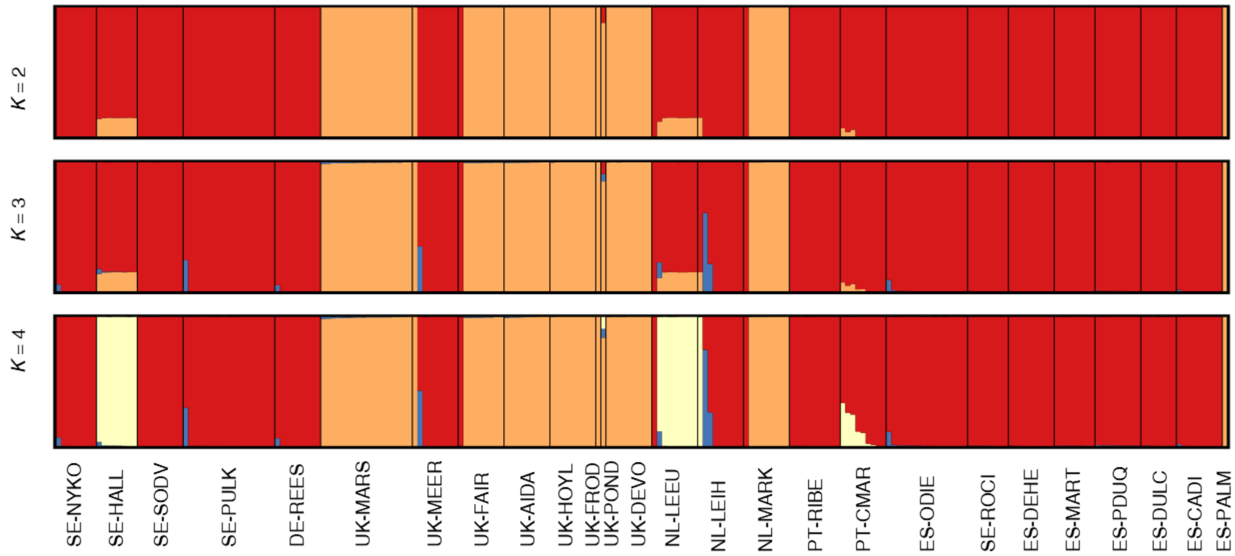


Fig. S3. Results of genetic assignments based on the Bayesian clustering analyses implemented in the program STRUCTURE (*K* = 2-4) for the dataset containing 230 individuals from Europe (5,692 SNPs). Each individual is represented by a vertical bar, which is partitioned into *K* coloured segments showing the individual's probability of belonging to the cluster with that colour. Thin vertical black lines separate individuals from different populations. Population codes as described in Table 1.

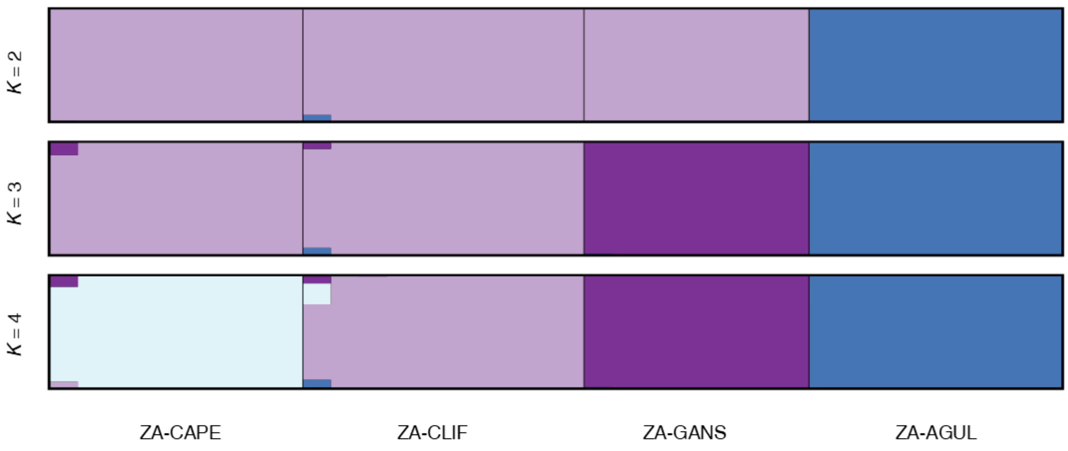


Fig. S4. Results of genetic assignments based on the Bayesian clustering analyses implemented in the program STRUCTURE (*K* = 2-4) for the dataset containing 36 individuals from South Africa (13,079 SNPs). Each individual is represented by a vertical bar, which is partitioned into *K* coloured segments showing the individual's probability of belonging to the cluster with that colour. Thin vertical black lines separate individuals from different populations. Population codes as described in Table 1.
